# A short nucleophosmin isoform couples nucleolar remodeling to DNA damage repair during meiosis

**DOI:** 10.64898/2026.11.10.765709

**Authors:** Sarah Potgieter, Saman Nayyab, Mia Uccello, Elizabeth M. Snyder

## Abstract

Nucleoli are nuclear compartments wherein ribosome biogenesis occurs. Observations spanning over a century have demonstrated that meiotic male germ cell nucleoli have unusual morphology and a close association with a large DNA damage repair foci formed around the sex chromosomes termed the XY-body. A mechanistic understanding of this relationship and its impact on ribosome biogenesis has remained elusive. High-resolution microscopy revealed XY-body associated nucleoli have an unusual two-part morphology made up of an XY-body intercalated and an XY-body adjacent part, each marked by different isoforms of the nucleolar protein nucleophosmin. Additionally, the entire structure is transcriptionally inactive and likely derived from autosomal nucleoli. Further, proper formation and maintenance of the structure rely on an interaction between NPM1 and the DNA damage marker H2AX. These findings provide mechanistic insight into the XY-body and “nucleolus” association and reveal unrecognized connections between nucleolar proteins, ribosome biogenesis, and DNA damage repair during meiosis.

## Introduction

The nucleolus is a subnuclear body that primarily functions in ribosome biogenesis (RiBi). In somatic cells, it is organized into three regions: the fibrillar center (FC), the dense fibrillar component (DFC), and the granular component (GC)^1^. In quiescent cells and mitotic cells in interphase, nucleoli are organized around sites of nascent rRNA transcription from ribosomal DNA repeats termed nucleolar organizing regions (NORs)^1^. Early in mitotic prophase, ribosome assembly is paused and the nucleolus disassembled to allow for even distribution of nucleolar components between daughter cells. In comparison, during meiosis in higher eukaryotes, nucleolar dissolution is delayed until the very end of prophase I^2–4^. Mammalian male germ cells are possibly the best example of this, with meiotic prophase I spanning twelve and a half days in mice and approximately twenty-four days in men^5^. How meiotic nucleoli are regulated during this unique developmental period remains unclear.

Previous work has demonstrated that the mammalian male meiotic nucleolus has unusual morphology^4,6–9^. Observations spanning over a century have demonstrated that the meiotic nucleolus is in close physical proximity to the XY-body^10–14^, a specialized nuclear region specific to male meiosis containing the sex chromosomes. Yet, in mice and men, neither the X nor Y contain an NOR^15,16^ thus the relevance or function of nucleolar association with the XY-body remains poorly understood.

The XY-body is a large nuclear domain enriched with DNA damage marks and repair factors that forms due to the inability of the X and Y chromosomes to fully synapse after meiotically programmed double-strand breaks (DSBs). As a result, sex chromosome DNA damage repair (DDR) is delayed relative to the autosomes, failing to be completed until just prior to meiosis II^17^. Recent reports have demonstrated pre-rRNA may be required for DDR and downstream meiotic sex chromosome inactivation (MSCI)^14,18–21^ suggesting there may be a functional relationship between XY-body-associated nucleoli and meiotic progression. However, no molecular basis for this relationship has been described.

In this study, we characterize the molecular and physical relationship between the meiotic nucleolus and the XY-body along with the activity of the nucleolus throughout meiotic prophase I. In doing so, we identify a poorly described isoform of the nucleolar protein nucleophosmin (NPM1) as a molecular link between the nucleolus, the XY-body, and DNA damage. These studies describe, for the first time since its discovery over a century ago, the drivers behind the formation of XY-body-associated nucleoli and reveal a previously unappreciated connection between sex chromosome DNA damage and nucleolar dynamics.

## Results

### The meiotic nucleolus has unusual morphology defined by distinct bodies

Previous molecular work describing the male meiotic nucleolus was hampered by use of isolated cells which alters nuclear architecture and lacks the developmental context necessary for precise prophase I substage determination^7,8,14,19,22^. Thus, there is no refined molecular description of *in situ* nucleolar morphology during meiosis and its relationship with the XY-body. To address this, we examined the localization of NPM1 (GC marker)^23^, fibrillarin (FBL, DFC marker)^24^, and nucleolin (NCL, a pan nucleolar marker)^25^ across male meiosis (Supp Fig. 1A). Somatic and mitotic testicular cells (Supp Fig. 1B) had classic nucleolar morphology with a region of FBL surrounded by a larger accumulation of NPM1 and NCL^26^.

Nucleolar organization in meiotic germ cells varied widely by meiotic phase (Fig. 1A). At zygotene, there was an accumulation of NPM1 that colocalized with neither FBL nor NCL. In pachytene spermatocytes a large, often non-spherical, <u>N</u>PM1-<u>F</u>BL-<u>N</u>CL region (NFN body) was observed near a much larger NPM1-only region (NPM1-only body) along with additional, smaller NFN bodies. By diplotene, most cells contained a single NFN body closely associated with a single NPM1-only body. After diplotene, NPM1-containing structures underwent dissolution with occasional small NPM1-only foci. This analysis reveals the GC marker NPM1 forms three morphologically distinct bodies during meiosis with developmental state-specific profiles: small and large NFN bodies and NPM1-only bodies. NFN bodies are presumably nucleoli while NPM1-only bodies represent a unique population of nuclear structures.

**Figure 1.**
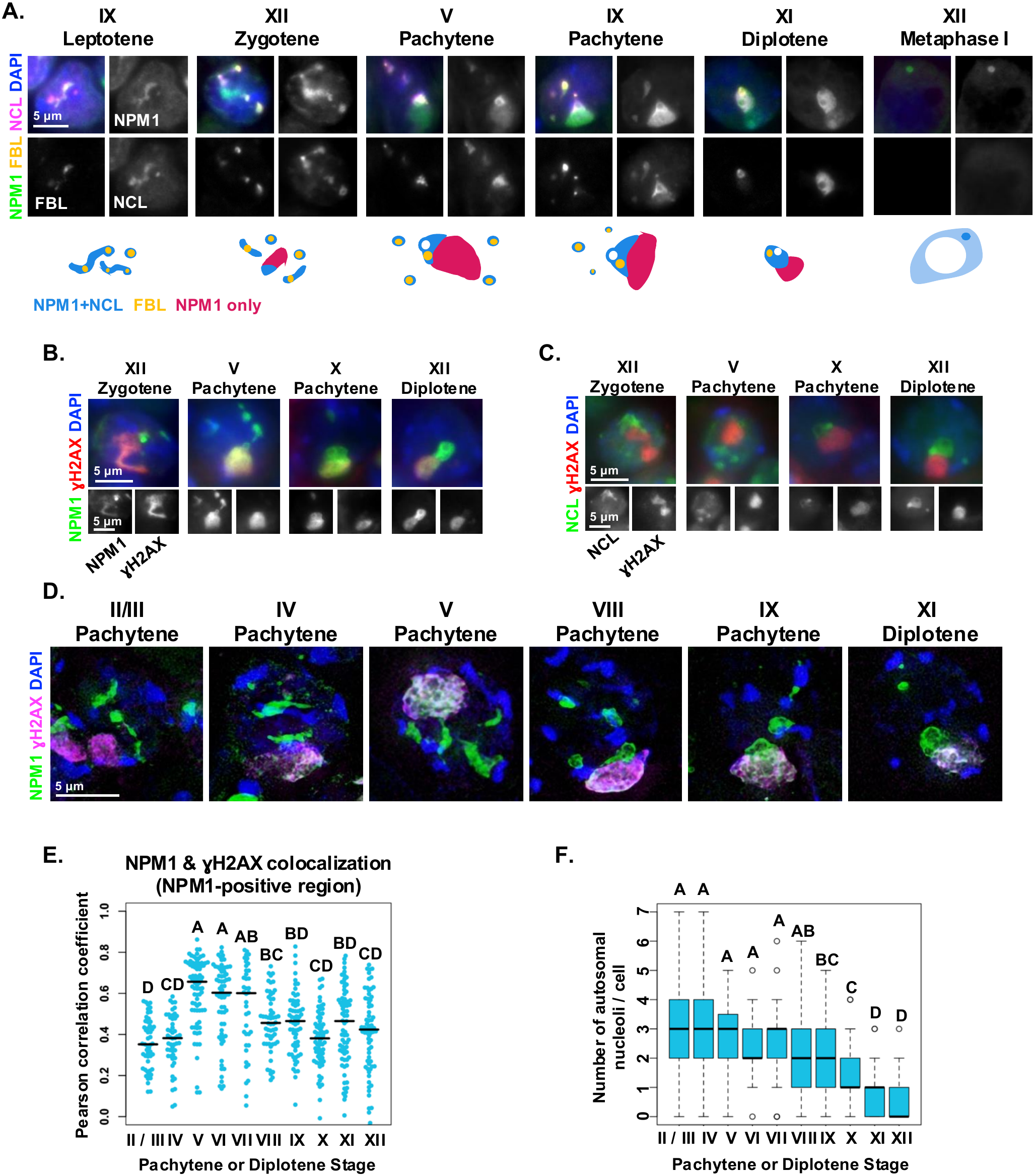
Two distinct NPM1-containing bodies differentially associate with the XY-body. Immunofluorescence of **(A)** NPM1, FBL, and NCL with graphical depiction of observed staining, **(B)** NPM1 and H2AX, and **(C)** NCL and H2AX in meiotic spermatocytes by developmental phase. **(D)** SIM standard deviation Z projections of NPM1 and H2AX in meiotic spermatocytes by developmental phase. **(E)** NPM1 and H2AX colocalization coefficients in XY-body associated NPM1 bodies by meiotic phase. Dots are individual cells (see Methods). **(F)** Number of non XY-body associated NPM1 bodies per cell by meiotic phase. (see Methods). Letters indicate significance levels determined by one-way ANOVA (p < 0.05). Bar – sample median. Values reported for individual cells from biological triplicate (14 - 23 cells / stage / animal). Images representative of biological triplicate. Roman numerals indicate spermatogenic stage.

### NPM1-only and NFN bodies form at the XY-body as a function of meiotic stage

Previous reports have suggested a close physical relationship between the nucleolus and the XY-body^8,10–14,19,27^. Given our identification of molecularly distinct NPM1 bodies throughout meiosis, we explored their relationship with the XY-body (Fig. 1B). A clear but imperfect colocalization of H2AX (a mark of the XY-body) and NPM1 but not NCL (Fig. 1C) was observed starting at zygotene, just prior to XY-body establishment^28^. Similar to previous observations^14^, H2AX completely colocalized with large NPM1-only bodies upon full XY-body formation at mid-pachytene^28^ and remained through diplotene. Additionally, H2AX was adjacent to a single, large NFN body (Fig. 1C). These analyses demonstrated both large NPM1-positive bodies associate with the XY-body in differential and developmentally regulated ways.

### NFN bodies are derived from XY-body and autosomal NPM1 sources

To better define the relationship between NPM1 bodies and the XY-body, we performed Structured Illumination Microscopy (SIM) (Fig. 1D) which demonstrated variable degrees of NPM1 intercalation within a network of H2AX throughout pachytene and diplotene. NPM1 and H2AX colocalization (Fig. 1E and Sup. Fig. 2A) was relatively low prior to XY-body formation (stage II/III – IV) but precipitously increased upon full XY-body maturation (stage V). As pachytene proceeded (stage VIII), an NPM1 body external to H2AX was observed coincident with overall reduced NPM1 and H2AX colocalization and formation of the XY-body-adjacent NFN body. Together, our observations demonstrate NPM1 is integrated within the XY-body and suggest that the adjacent NFN body may be partially derived from NPM1 within the XY-body.

We next asked if XY-body-adjacent NFN bodies could be further derived from autosomal NFN bodies as occurs in somatic cells wherein small nucleoli fuse to generate larger ones following mitotic division^29^. Quantification of autosomal NPM1 revealed a significant decrease in the number (Fig. 1F) and average volume (Sup Fig. 2B) of non-H2AX-adjacent NPM1-marked bodies simultaneous to formation of the XY-body-adjacent NFN body despite constant autosomal NPM1 pixel intensity (Supp Fig. 2C), supporting the notion that XY-body-adjacent NFN bodies are additionally derived from autosomal sources. In sum, our imaging analyses demonstrated 1) NPM1-only bodies form at nascent XY-bodies, 2) NFN bodies are likely derived in part from XY-body-intercalated NPM1, and 3) as meiosis progresses NFN bodies further accumulate nucleolar components from autosomal nucleoli.

### NFN and NPM1-only bodies contain different resident NPM1 isoforms

NPM1 is observed as three distinct protein isoforms (Fig. 2A) which display different nucleolar localization preferences^30^ and are encoded by three alternative mRNAs (Fig. 2B). Thus, we aimed to determine which NPM1 protein isoforms were resident to NFN and NPM1-only bodies. Isoform-specific RT-PCR (Fig. 2B-C) demonstrated two NPM1 isoforms were detectible in adult testis, one encoding the full length NPM1 (NPM1-L) and a shorter isoform encoding a truncated peptide with an alternative C-terminus (NPM1-S). Western blot using the same polyclonal anti-NPM1 antibody as in Fig. 1 across testicular development (Fig. 2D) demonstrated two bands of the expected molecular weights for NPM1-L and NPM1-S. Those same samples were then probed with a monoclonal anti-NPM1 antibody (Fig. 2D) which detected a single band. Dual-labeled western blotting with both antibodies (Fig. 2E) confirmed monoclonal anti-NPM1 recognized a peptide corresponding to NPM1-L while polyclonal anti-NPM1 (herein referred to as panNPM1) recognized two peptides corresponding to NPM1-L and NPM1-S. Isoform expression profiles using tissue-level RNA-sequencing data^31,32^ (Supp Fig. 3A) demonstrated reproductive tissues expressed higher levels of the short NPM1 encoding mRNA, an observation confirmed on the protein level (Supp Fig. 3B).

**Figure 2.**
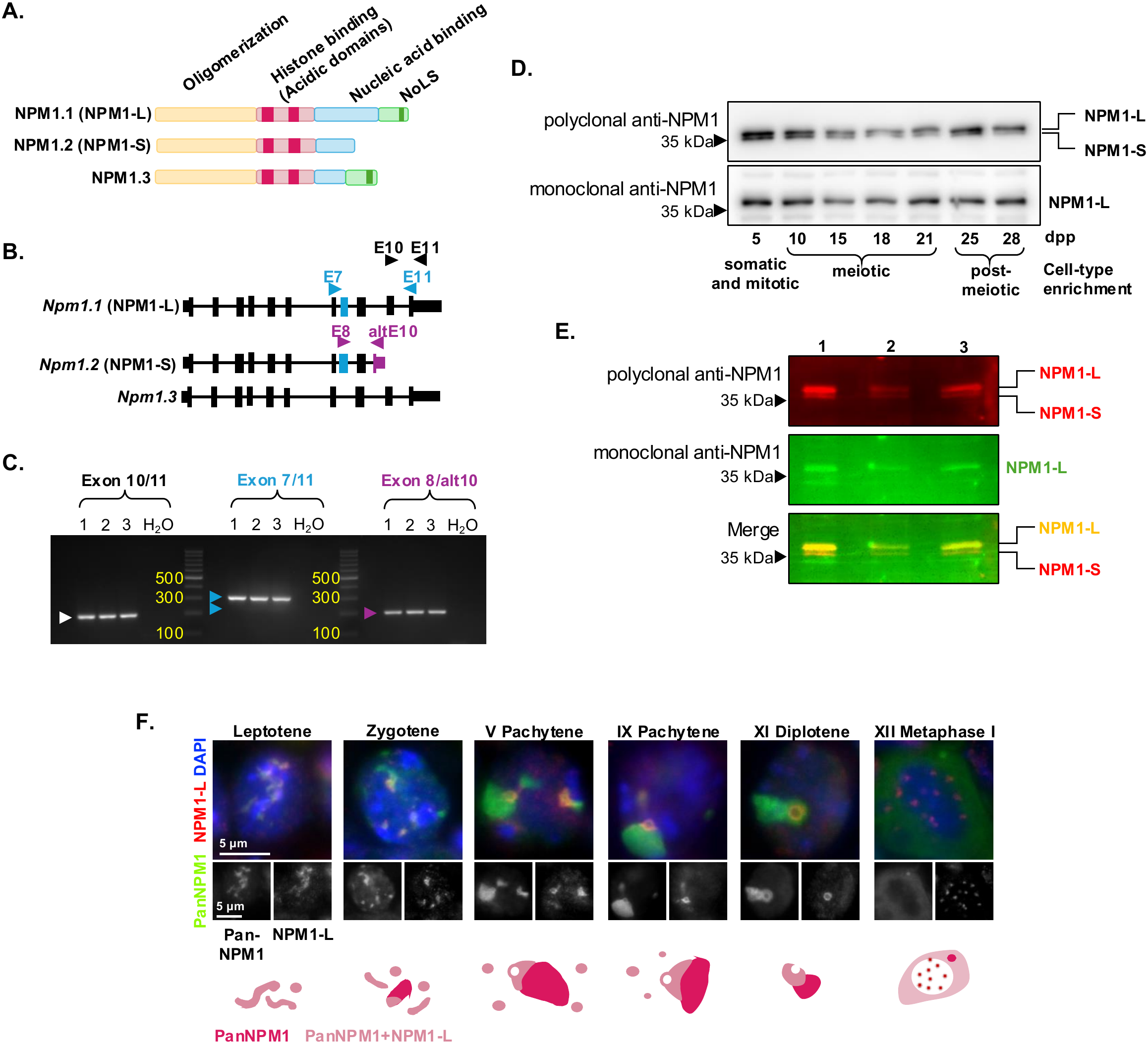
Distinct NPM1-containing bodies in meiotic male germ cells are defined by distinct resident NPM1 protein isoforms. Schematics of **(A)** NPM1 protein isoforms and **(B)** *Npm1* mRNA isoforms and RT-PCR detection approach. Arrowheads – priming sites. Plum – *Npm1.2*-specific exon and diagnostic primers (expected product: 202bp). Blue – *Npm1.3*-excluded exon and diagnostic primers (expected products – *Npm1.1*: 319bp, *Npm1.3*: 234bp). **(C)** RT-PCR of testicular *Npm1* isoforms. Numbers – biological replicates. Arrowheads – expected products as in (B). **(D)** Western blot of two NPM1 antibodies on whole-testis lysate across development. Blots representative of biological triplicate. dpp – days post-partum. **(E)** Dual western blot of NPM1 antibodies in (D) on adult whole-testis lysate. Numbers – biological replicates. **(F)** Immunofluorescence and graphical model of polyclonal (panNPM1) and monoclonal (NPM1-L) NPM1 antibody staining in meiotic spermatocytes by developmental phase. Images representative of biological triplicate. Roman numerals indicate spermatogenic stage. See Sup Fig. 7 for western blot loading controls.

Leveraging the ability to distinguish between the two NPM1 isoforms, we next asked whether they corresponded to different NPM1-containing bodies within meiotic germ cells (Fig. 2F). As NPM1-only bodies formed during zygotene, NPM1-L was observed primarily in small, discrete foci or as a singular non-spherical body similar to small and large NFN bodies, respectively, while panNPM1 detected those same foci along with larger regions morphologically identical to NPM1-only bodies. Thus, the two NPM1 bodies are distinguished by their relative NPM1 protein isoform abundance with NFN bodies containing either long or both NPM1 isoforms and NPM1-only bodies marked primarily by the short isoform.

### Neither NPM1-only nor XY-body adjacent NFN bodies are true nucleoli

NPM1-only and XY-body-adjacent NFN bodies are neither spherical nor contain centrally embedded FBL granules, leading us to wonder whether either function as normal nucleoli. We first asked whether either were analogous to a nucleolar cap, a large condensate formed when nucleolar function is inhibited^33^, by localization of PTB-associated splicing factor (PSF) which relocates to the nucleolus during cap formation^33^. Throughout meiosis, PSF remained nucleoplasmic and was excluded from NPM1-positive regions (Supp Fig. 4A-B) suggesting XY-body-associated NPM1-containing bodies are not nucleolar caps. Supporting this notion, ribosomal proteins RPL7A and RPL26 / 26L1 were also observed in all NFN bodies throughout meiosis (Supp Fig. 4C-D).

The definitive measure of an active nucleolus is expression of rDNA^34^. Early studies of rDNA expression during meiosis suggested pre-rRNA abundance peaked in mid-pachytene followed by a decrease around late pachytene^7,35,36^. More recent reports suggest nascent pre-rRNA levels ebb across meiosis^14^. To better refine this and determine pre-rRNA localization relative to the newly defined NPM1 bodies, we examined the localization of 47S pre-rRNA by RNA-FISH relative to NPM1 and H2AX (Fig. 3A). High 47S rRNA signal indicative of an active nucleolus was detected in most NPM1-positive bodies in early meiosis before decreasing through pachytene with only weak signal remaining at late diplotene suggesting that RiBi correlates with autosomal NFN body decline. Additionally, all NPM1-positive bodies associated with H2AX were 47S rRNA negative. Thus, despite the inclusion of ribosomal proteins and the exclusion of nucleolar cap proteins, neither XY-body-adjacent NFN nor NPM1-only bodies are transcriptionally active nucleoli.

**Figure 3.**
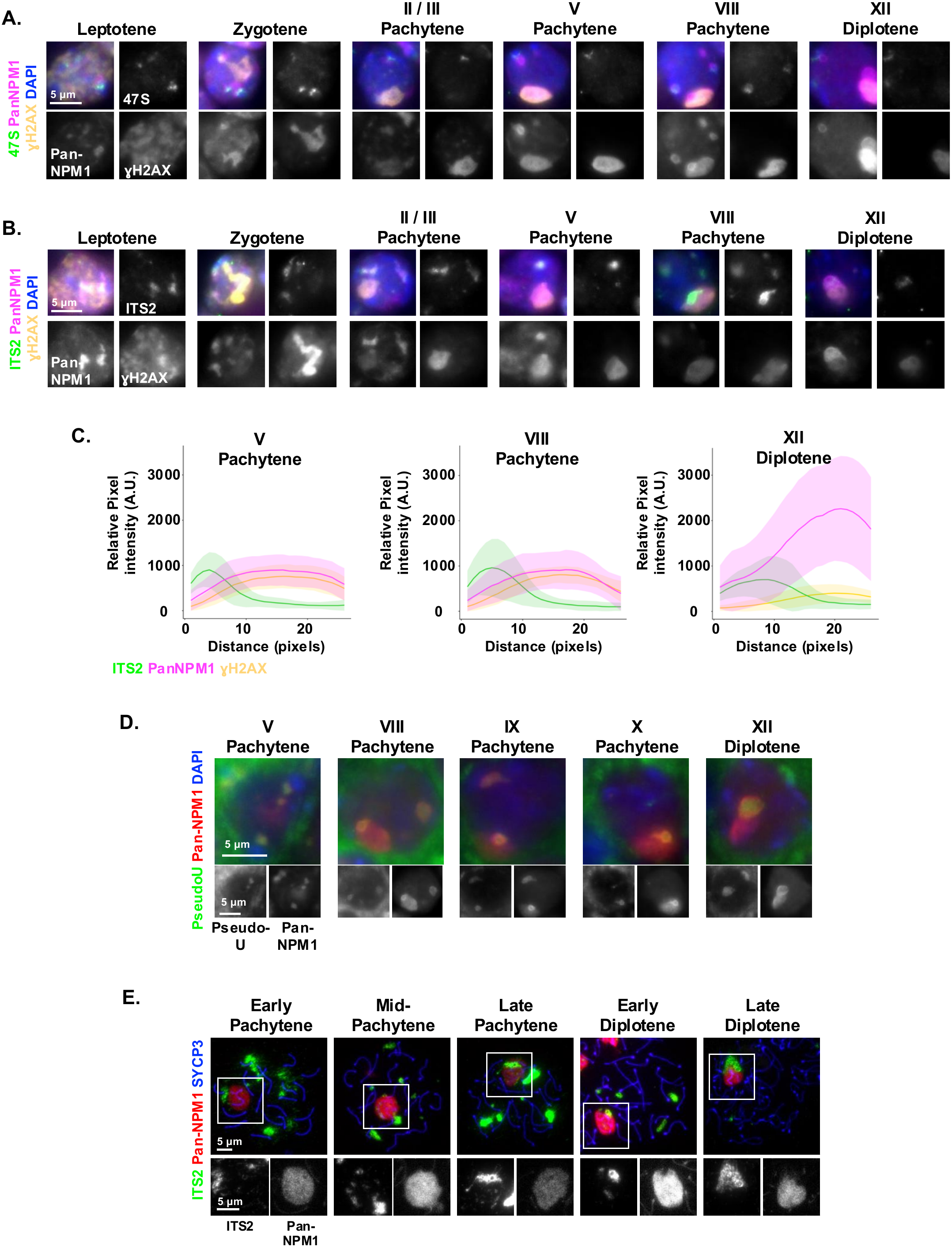
XY-body intercalated NPM1-only bodies do not contain rRNA though NFN bodies do. Immunofluorescence of panNPM1 and H2AX in meiotic spermatocytes by development phase combined with RNA-FISH for **(A)** the 5’ ETS sequence of 47S pre-rRNA and **(B)** ITS2 containing pre-rRNA. **(C)** Plot profile of ITS2 pre-rRNA, panNPM1, and H2AX across both XY associated NPM1-containing bodies. Solid line – averaged pixel intensity. Shade – standard deviation. Values reported for individual cells from biological triplicate (8 – 20 cells / stage / animal). **(D)** Immunofluorescence of panNPM1 and pseudouridine (pseudoU) in meiotic spermatocytes by development phase. **(E)** RNA-FISH of ITS2 pre-rRNA with immunocytochemistry of SYCP3 and pan NPM1 on spermatocyte nuclear spreads. Images representative of biological triplicate. Roman numerals indicate spermatogenic stage.

### XY-body-adjacent NFN but not NPM1-only bodies contain pre-rRNAs

Previous reports suggest pre-rRNA may be required for DDR and MSCI based in part on observations of pre-rRNA within the XY-body^14,19^. To determine if we observed a similar phenomenon and relate it to our newly defined NPM1 bodies, we localized partially processed pre-rRNA containing the internal transcribed spacer two (ITS2) relative to NPM1 and H2AX across meiosis I (Fig. 3B-C). Unexpectedly, we failed to observe ITS2 colocalized with H2AX regardless of meiotic stage, though ITS2 was observed adjacent to the large H2AX regions of pachytene and diplotene cells. These observations support our previous supposition that XY-body-associated NFN bodies are derived, in part, from autosomal nucleoli which are the presumed originators of partially processed pre-rRNA. To determine whether any rRNA was detectible in the XY-body, we examined the localization of pseudouridine (Fig. 3D), an abundant rRNA modification deposited during rDNA expression^37,38^. Although localized to all NFN bodies, pseudouridine was consistently excluded from NPM1-only bodies and therefore XY-bodies as well.

### XY-body-associated NPM1-positive bodies are sensitive to analysis approach

The above analyses demonstrate that, contrary to previous reports^14,19^, the XY-body does not contain pre-rRNA. To better understand this disconnect, we critically evaluated the impact of nuclear spreads on the architecture of the two XY-body-associated NPM1-positive bodies. Upon formation of the XY-body and throughout the remainder of development, panNPM1 was observed throughout the γH2AX-positive region (Sup Fig. 5A), which was occasionally accompanied by a small, discrete foci of additional NPM1 signal.

Given this, we assessed NFN bodies in nuclear spreads by examining NCL localization (Sup Fig. 5B). In striking contrast to intact tissue observations, small foci of NCL appeared embedded in the large NPM1 region. We observed a similar “mixing” of the two NPM1 bodies in nuclear spreads stained for both NPM1-L and panNPM1 (Sup Fig. 5C) suggesting the separation between NPM1-only and XY-body-adjacent NFN bodies was not maintained during nuclear spread processing.

To determine if collapsed NPM1-containing bodies during sample processing could explain observations of pre-rRNA in the XY-body, we stained nuclear spreads for ITS2 and either panNPM1 (Fig. 3E) or γH2AX (Supp Fig. 5D). In many cells, especially in mid- to late pachytene, ITS2-positive foci were within either the large panNPM1 or γH2AX signal, mirroring the pattern of NCL. These analyses confirmed a lack of pre-rRNA in the XY-body and further demonstrated nuclear spreads artificially mix the two NPM1-containing bodies at specific developmental stages. Together, these observations explain why pre-rRNA has previously been detected in the XY-body and demonstrate why the two distinct NPM1 bodies went undetected in previous studies, which have largely been limited to nuclear spread analyses^4,11,14,19,22^.

### NPM1 interacts with□H2AX

Our earlier analyses demonstrated NPM1-S directly localized to the XY-body. Previous work has demonstrated NPM1 localization at ionizing radiation-induced foci^39^. Given this, we investigated whether the localization of NPM1-S to the H2AX-positive XY-body could be mediated by their interaction. Immunoprecipitation with panNPM1 in adult testes probed for NPM1 and□H2AX (Fig. 4A) robustly detected NPM1-S and H2AX, revealing that NPM1 and H2AX interact suggesting NPM1-S accumulation within the XY-body and subsequent formation of the adjacent NFN body may be mediated by NPM1-S’s affinity for□H2AX.

**Figure 4.**
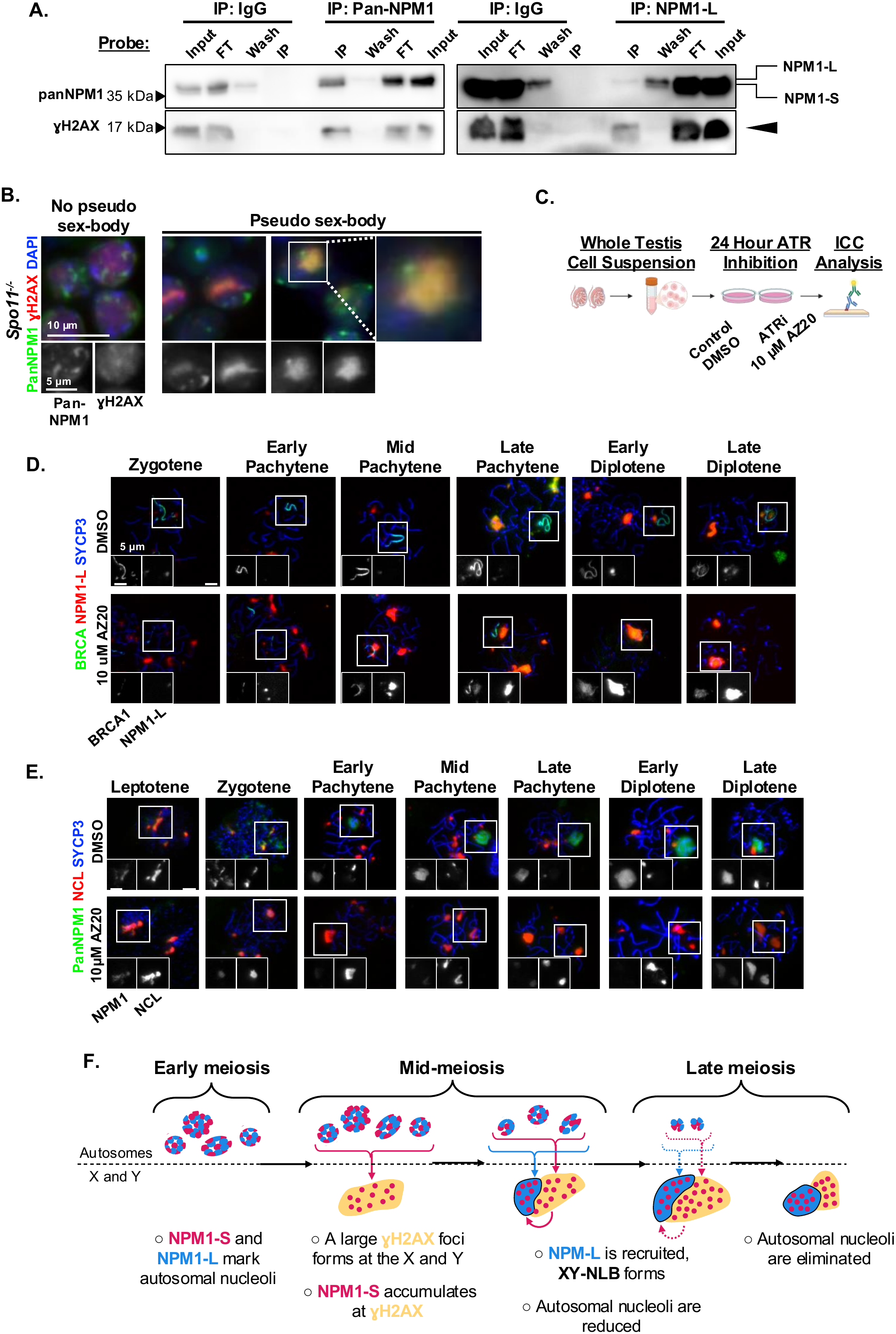
XY-body associated NPM1-containing bodies rely on formation and maintenance of a H2AX domain. **(A)** Immunoprecipitation of NPM1 by pan- or NPM1-L specific antibodies with IgG isotype controls in adult whole-testis lysates. Blots representative of biological triplicate. Input – total testis lysate, FT – flow-through, Wash – first bead wash post binding, IP – immunoprecipitate. Arrowhead - H2AX. Both H2AX blots have been horizontally flipped to match the loading order of the NPM1 blots (**B)** Immunofluorescence of panNPM1 and H2AX in *Spo11* mutant spermatocytes with or without a pseudo-sex body. **(C)** Primary spermatocyte ATR inhibition cultures. ICC – immunocytochemistry. Generated via BioRender. ICC of **(D)** BRCA1, NPM1-L, and SYCP3 and **(E)** panNPM1, NCL, and SYCP3 on cultured spermatocytes. **(F)** Model of nucleolar association with the XY-body throughout meiosis. Images representative of biological triplicate. See Sup Fig. 8 for western blot loading controls.

To further test this supposition, we asked whether formation of NPM1-only bodies was specific to the XY-body or to large□H2AX-positive foci in general. To do so, we leveraged *Spo11* knockout (*Spo11^-/-^*) testes, which contain occasional meiotic cells that accumulate large regions of□H2AX independent of DSBs^40^ and the sex chromosomes^41^, termed pseudo sex-bodies^41,42^. Immunofluorescence of panNPM1□and H2AX (Fig. 4B) demonstrated NPM1 localization similar to that observed in leptotenes in *Spo11^-/-^* cells without a pseudo sex-body. In contrast, the pseudo sex-bodies of *Spo11^-/-^* cells associated with NPM1-marked foci reminiscent of pachytene NPM1-only and NFN bodies, a molecular designation confirmed by their NCL localization (Supp Fig. 6A). Together, these analyses suggest formation of both NPM1-only and XY-body-adjacent NFN bodies relies on accumulation of H2AX and support the notion that NPM1-S accumulation at the XY-body is a function of H2AX affinity.

### NPM1-only foci are reliant on sustained deposition of γH2AX

Our results reveal both NPM1-only and NFN bodies form at large accumulations of H2AX and that NPM1 interacts with H2AX. To determine whether H2AX deposition impacts NPM1 association with the XY-body, we cultured primary testicular cells with AZ20^43^ (Fig. 4C), an inhibitor of Ataxia Telangiectasia and Rad3-related protein (ATR) which drives H2AX phosphorylation in the XY-body^44,45^. Twenty-four hours of AZ20 treatment led to a complete loss of XY-body H2AX (Supp Fig. 6B) in primary spermatocytes. Using this model, we assessed the impact on XY-body-associated NFN bodies by co-staining treated spermatocytes with NPM1-L and BRCA1, a DDR protein that marks the sex chromosome axes^46^ independent of H2AX^43^ (Fig. 4D). Untreated spermatocytes contained small, sex-chromosome-adjacent NPM1-L foci starting in mid-pachytene followed by partial colocalization of BRCA1 and NPM1-L as meiosis progressed to diplotene. In contrast, AZ20 treatment resulted in very large NPM1-L positive foci near axial BRCA1 in mid-pachytene spermatocytes. These became more prominent with abundant accumulation of BRCA1, presumably from the axes, in later spermatocytes.

The profound impact of AZ20 treatment on NFN bodies (marked by NCL) led us to next ask whether NPM1-only bodies were similarly affected (Fig. 4E). In untreated spermatocytes, NPM1 initially localized with NCL followed by localization to both NFN and NPM1-only bodies as observed in testicular spermatocytes. Following AZ20 treatment, NPM1 was greatly reduced in early spermatocytes similar to recent reports^14^. Virtually no NPM1 was detected in later spermatocytes in contrast to dramatic increases in NPM1-L under the same conditions. These observations first suggest panNPM1 preferentially stains NPM1-S in spermatocytes spreads and further demonstrates NPM1-S is uniquely sensitive to H2AX loss, with NPM1-S accumulation and thus formation and maintenance of NPM1-only bodies reliant on normal H2AX deposition. Together, analyses of NFN and NPM1-only bodies show that both XY-body-associated NPM1 bodies are sensitive to H2AX and suggest a connection between NPM1 and DDR via BRCA1 in meiotic male germ cells.

## Discussion

The physical relationship between the nucleolus and the XY-body of meiotic male germ cells was reported more than a century ago^47^. This work is the first to describe the molecular mechanisms underpinning their relationship. Our observations reveal a two-part structure composed of an XY-body-intercalated element (NPM1-only bodies) and an adjacent nucleolus-like body (NFN bodies), each marked by a different protein isoform of NPM1. Neither body is an active nucleolus by classic definition. Specifically, we identify a poorly studied C-terminus truncated NPM1 isoform as the major NPM1 component of the XY-body and a likely driver of the reported nucleolus-XY-body association. We further identify the interaction between NPM1 and the DNA damage mark H2AX as a prerequisite for proper formation and maintenance of the nucleolus-XY-body association. Based on our observations, we propose a model (Fig. 4F) whereby NPM1-S’s affinity for H2AX drives NPM1 accumulation within a newly forming XY-body. Subsequently, NPM1-S separates from the XY-body and recruits NPM1-L to form an XY-body-adjacent <u>n</u>ucleolus-like <u>b</u>ody (XY-NLB). As meiosis progresses, the XY-NLB accumulates nucleolar components from the XY-body and active autosomal nucleoli, driving reduction and eventual loss of autosomal nucleoli. These observations provide insight into the mechanisms of XY-body and “nucleolus” association and identify the short form of NPM1 as an important player in nucleolar regulation. They further reveal previously unappreciated connections between DDR, nucleoli, and RiBi during meiosis.

Although this work established the mechanism driving the nucleolus-XY-body association, their direct relationship with RiBi is not entirely defined. Contrary to previous suppositions^4,8,11,12,19,27,35^ which assumed XY-NLBs were functional nucleoli, we find them to be transcriptionally inactive. Based on partially processed and modified rRNAs and ribosomal proteins localized to XY-NLBs, it is feasible they maintain partial nucleolar function wherein nascent rDNA expression is absent but other aspects of RiBi continue. Supporting this notion, XY-NLB accumulation of nucleolar components at the apparent cost of autosomal nucleoli suggests they modulate the eventual shut down of RiBi at the end of the meiotic cell cycle. Alternatively, they may serve as a storage site for RiBi components to be activated after completion of meiosis, analogous to nucleolar protein enriched perichromosomal regions observed during mitosis^48^.

In contrast, the function of NPM1-only bodies is unlikely to be RiBi related. However, nucleolar proteins have been implicated as regulators of DDR and MSCI^14,39,49–51^ suggesting their close association with the XY-body may be a function of their role therein. NPM1, in particular, has been linked to DDR in multiple systems^52,53^ wherein NPM1 localizes to sites marked by H2AX upon DNA damage^39^ and facilitates repair, likely through interactions with BRCA1^54^. Unfortunately, most published studies focus on full length NPM1 or fail to distinguish between protein isoforms. Based on our observation of distinct NPM1 protein isoforms differentially associating with the XY-body, it seems feasible that NPM1-S may have a specific role in DDR during meiosis. This assertion is further supported by our observation that NPM1-S is enriched in reproductive tissues, appears to have high affinity for H2AX, and is sensitive to H2AX loss despite increases in NPM1-L and other nucleolar proteins. A more refined understanding of NPM1-S and its protein interactions during DDR may reveal important aspects of DDR regulation.

A confounding observation is the lack of pre-rRNA in the XY-body, which has been reported to contain a wide range of partially processed rRNAs presumed to be important for DDR^14,19^. It is possible we fail to detect pre-rRNAs or their modifications within the XY-body due to assay sensitivity. Alternatively, and more likely, detection of pre-rRNAs in previous work results from perturbations of XY-body-associated NPM1-containing bodies during sample processing. Nuclear spreads are a common approach to study subnuclear localization in meiotic germ cells^55^, yet they require chemistries known to negatively impact phase separated condensates such as altered pH and tonicity^56–62^. The striking collapse of separated NPM1 bodies observed in nuclear spreads highlights the need for careful assay selection when studying the nucleolus and DDR foci.

While RiBi and DDR are well described functions for the nucleolus and/or nucleolar proteins such as NPM1, alternative functions are also possible. For example, XY-NLBs bear a striking resemblance to another nucleolus-like body observed in oocytes just prior to fertilization. Oocyte NLBs were originally presumed to store nucleolar proteins to support RiBi after fertilization^63^. However, recent reports have suggested oocyte NLBs act to regulate chromatin state of parental genomes^64^. Given spermatocyte XY-NLB’s proposed role in MSCI^14^, it is possible the two NLBs are analogous. Whether male germ cell XY-NLBs influence post-meiotic male germ cell chromatin remains an open question.

The work described here defines the structure and molecular composition of poorly described, but long known, nucleolus-like bodies in meiotic male germ cells. These studies reveal a previously undescribed mechanism regulating RiBi at the end of meiosis and further identify NPM1-S as a potential new player regulating sex chromosome DDR. Although these studies do not define the entirety of XY-body-associated NPM1-S function, they lay the foundation for further studies focused on RiBi regulation in male germ cells and NPM1-S’s specific function in DDR.

## Resource Availability

### Lead contact

Requests for further information and resources may be directed to and will be fulfilled by the lead contact, Elizabeth Snyder

### Materials availability

This study did not generate new unique reagents.

### Data and code availability

This study does not report original code.

This paper analyzes existing, publicly available data accessible at The Encode Project^31,32^ (accession numbers ENCFF286WTQ, ENCFF358NPU, ENCFF445AWP, ENCFF958CHE, ENCFF871XHK, ENCFF952JKH, ENCFF104UFH, ENCFF126WYO, ENCFF517RDO, ENCFF786ZKB, ENCFF682XSC, ENCFF690HKC, ENCFF492PRP, ENCFF581OEV, ENCFF161LEK, ENCFF516HOO, ENCFF312OKA, and ENCFF463WEH).

Any additional information required to reanalyze the data reported in this paper is available from the lead contact upon request.

## Supporting information

RNA-FISH Probe Sequences

## Acknowledgements

The authors would like to thank Yeva Shamailova, Megan Gilmore, and Ashley Di Agostino for support with animal husbandry; the Waksman Institute Shared Imaging Facility, Rutgers, The State University of New Jersey for their support in SIM imaging, Drs. Devanshi Jain and Ignasi Roig Navarro for the gift of *Spo11* mutant testis, Dr. Satoshi Namekawa for the gift of the BRCA1 antibody, and Dr. Mary Ann Handel for critical evaluation and intellectual contribution throughout project development. The authors would especially like to thank our funding sources; the Eunice Kennedy Shriver National Institute of Child Health and Human Development (NIH-NICHD R03 HD108418 and R01 HD107066 to ES along with F31 HD116499 to SP).

## Author Contributions

SP – methodology, validation, investigation, data curation, writing, visualization; SN – methodology, investigation; MU – methodology, investigation; ES – conceptualization, methodology, validation, formal analysis, investigation, resources, data curation, writing, visualization, supervision, project administration, funding acquisition.

## Supplemental Information titles and legends

Document S1: Supplemental Figures 1-8

Document S2: Supplemental Figure Legends

Document S3: RNA-FISH Probe Sequences

Supplementary Table 1: Antibody Conditions

## Star Methods

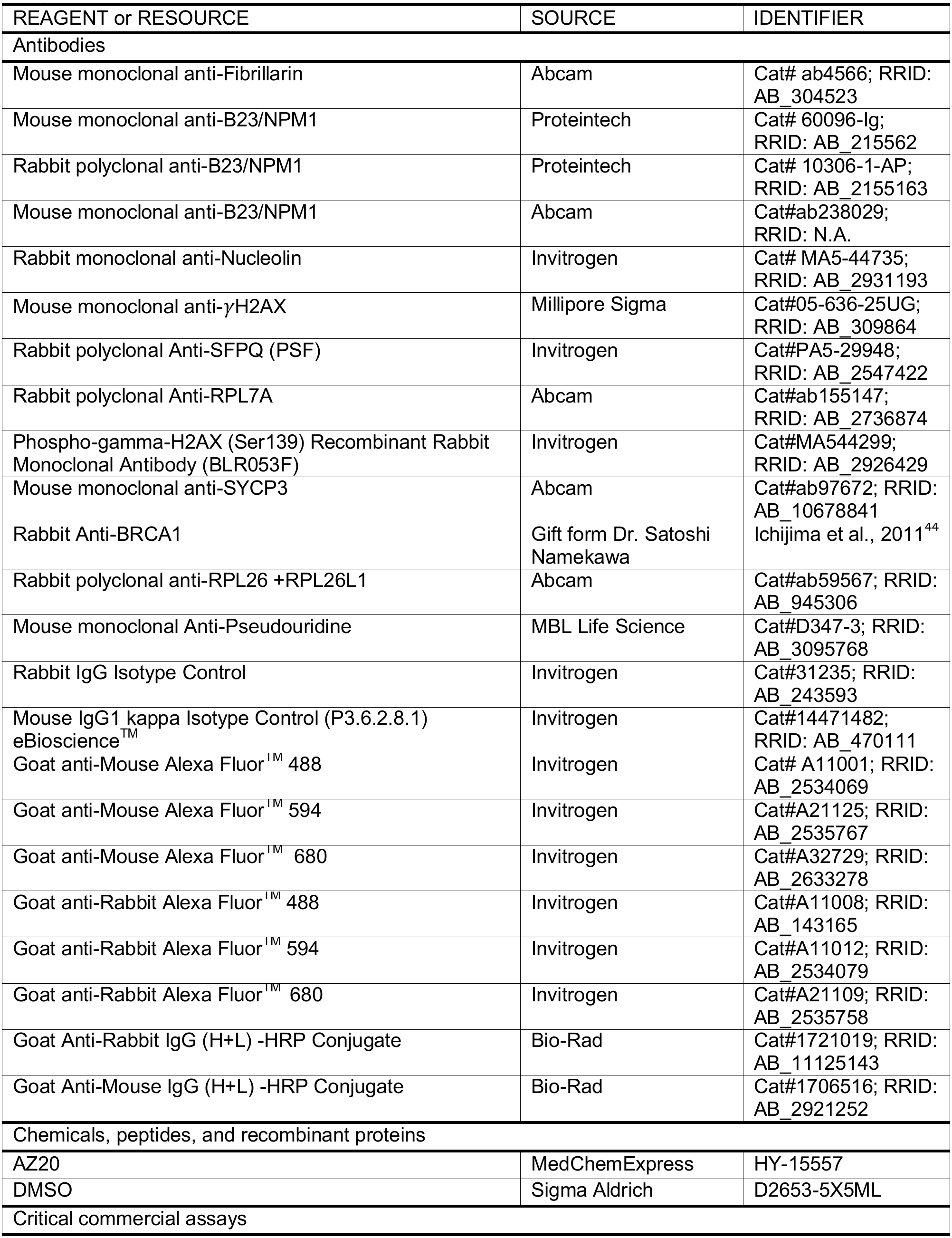
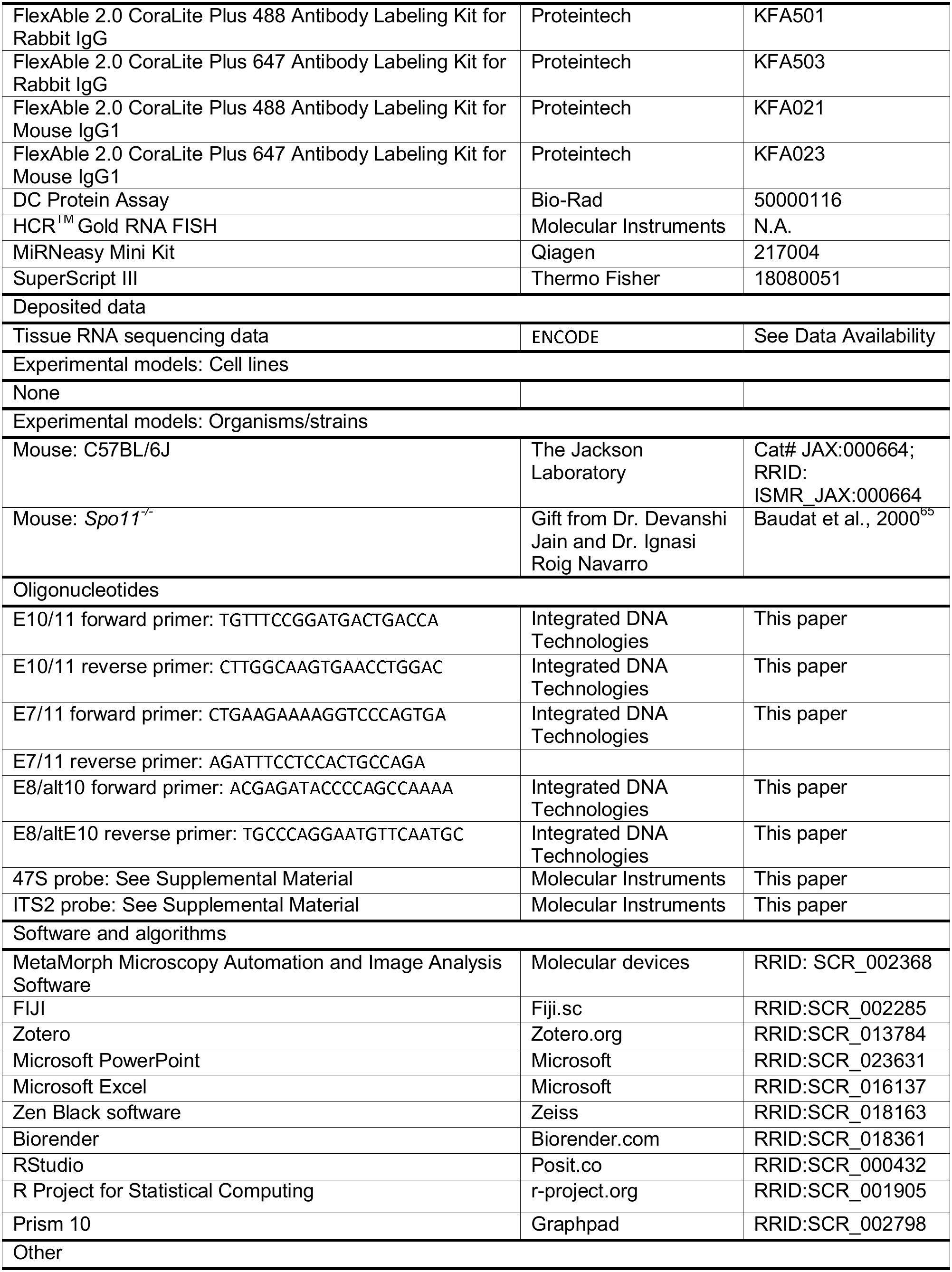
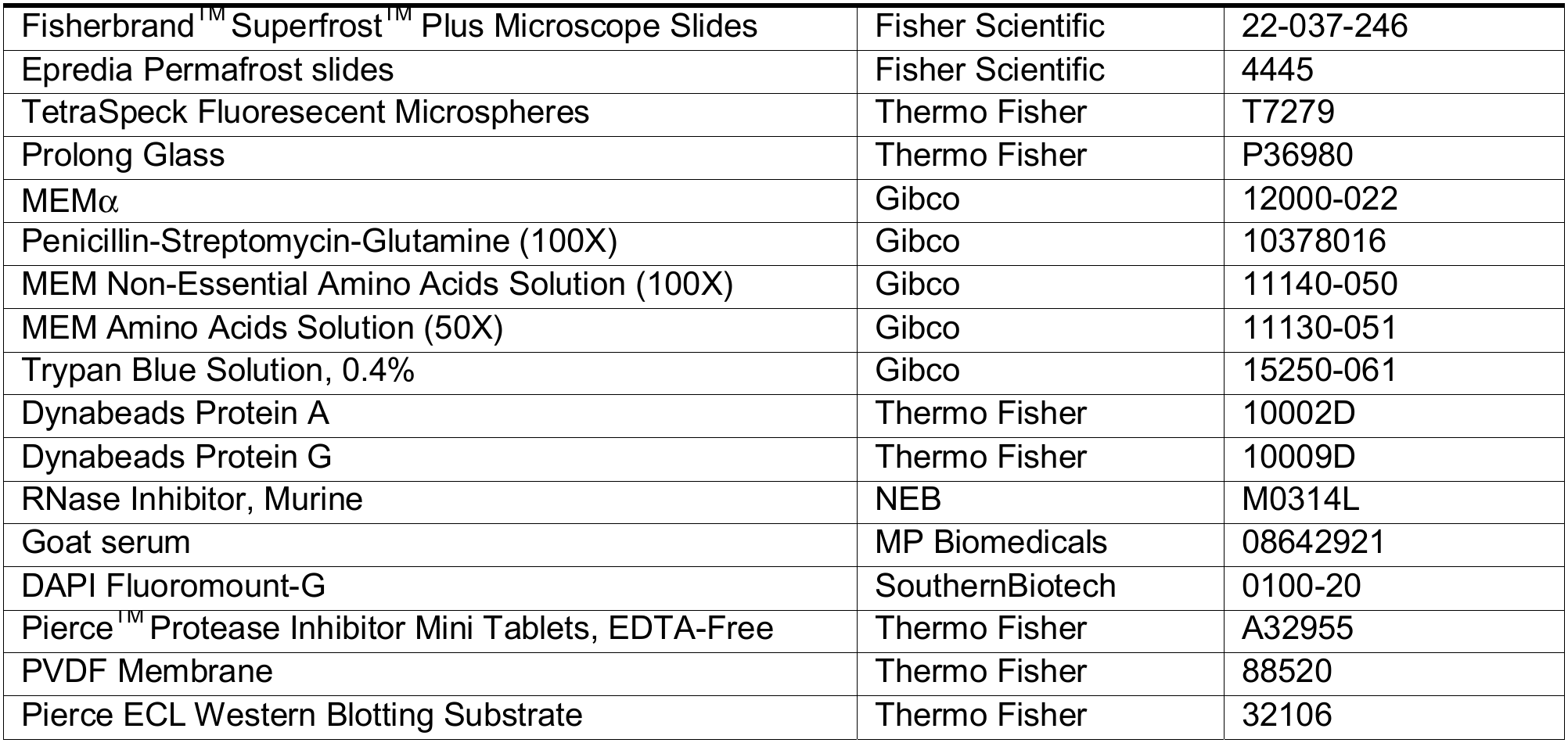
Key Resources Table.

## Experimental model and study participant details

### Mice

Founder C57BL/6J mice were purchased from The Jackson Laboratory and used to establish an in-house breeding colony which was refreshed every six to nine months with newly purchased C57BL/6J females. All experiments were performed using mice aged 60-180 dpp unless otherwise indicated. All protocols involving animal care and husbandry were approved and performed in accordance with the guidelines from the Rutgers Institutional Animal Care and Use Committee and were within AALAC and IACUC guidelines. Mice were kept in sterile, climate-controlled conditions, with a consistent 12 - h light - dark cycle. Mice were fed irradiated rodent chow (LabDiet 5058) and had access to food and water ad libitum.

## Method details

### Immunofluorescence

C57BL/6J testes were collected and fixed in 4% (w/v) PFA for 20 hours at 4°C. Tissues were subsequently rinsed in PBS, dehydrated in ethanol, embedded in paraffin wax, and cut into 4 μm sections. *Spo11-/-* samples were generous gifts from Drs. Devanshi Jain and Ignasi Roig Navarro. Immunofluorescence was done as described in^66^. In cases where antibodies derived from the same species were multiplexed, primary antibody was directly labeled using FlexAble 2.0 CoraLite Plus Antibody Labeling Kits (Proteintech) according to the manufacturer’s instructions. For antibody condition details, see Supplementary Table 1. Slides were visualized as described below. A biological replicate of *n = 3* was used for all quantification and visualization. The stage of the cycle of the seminiferous tubules was determined based on parameters described in^66,67^.

### Preparation of nuclear spreads and immunocytochemistry

C57BL/6J testes from 60 - 180 dpp animals were collect in PBS and nuclear spreads followed by immunocytochemistry was performed as in^68^. For antibody conditions, see Supplementary Table 1. Spermatocyte stage was determined by SYCP3 staining pattern as described previously^68^. A biological replicate of *n = 3* was used for all visualization and quantification.

### Preparation of cell cultures and immunocytochemistry

C57BL/6J testes from 60 - 180 dpp animals were collect in Enriched Krebs-Ringer Bicarbonate medium (120 mM NaCl, 4.8 mM KCl, 25.2 mM NaHCO_3_, 1.2 mM KH_2_PO_4_, 1.2 mM MgSO_4_.7H_2_O, 1.3 mM CaCl_2_, 1X Penicillin-Streptomycin-Glutamine (Gibco), 1X essential amino acid (Gibco), 1X nonessential amino acid (Gibco), 11.1 mM Dextrose) and a single cell suspension prepared for cell culture as described in^43^. Cell number was assessed using Trypan blue (Gibco) for a final dilution of 2.5 x 10^6^ cells / mL in MEMα (Gibco) containing either DMSO (Sigma, control media) or ATR inhibitor (MedChemExpress, 10 μM) and cultured for 24 hours at 32°C with 5% CO_2_ as in^43^. After culture, viability was assessed again using Trypan blue, with only clear cells counted as viable. Cultured cells were then processed for nuclear spreads and immunocytochemistry was performed as in^43^.

### Microscope imaging

Standard microscopy images were captured on a custom-built Zeiss microscope with both brightfield and fluorescent capabilities using MetaMorph imaging software (molecular devices) version 7.10.5.476. Each channel was individually imaged, and the channels were color-combined using either the built-in color combine tool or FIJI (Fiji.sc) in the case of quadruple channels. Within an experiment, each channel was captured using a consistent exposure time and signal intensity was matched across images by either matching background signal intensity or by setting the same min and max values on the pixel intensity histogram. All images are representative of a biological triplicate or greater.

### Structured Illumination Microscopy (SIM) and data processing

Images were captured on a Zeiss Elyra7 Structured Illumination with Lattice SIM^2^, with Ziess Axio Observer 7 microscope base, Piezo X-Y stage and Z-Piezo. The numerical aperture of the objective was 63x oil / 1.4 NA and dual sCMOS PCO Edge 4.2, 2048×2048 format, 6.5μm pixel size cameras were used. 13 Lattice illumination phases were used, and the Z step size was 0.13μm. To ensure alignment of all channels, a Z stack of 0.1μm TetraSpeck beads (Thermo Fisher) mounted in Prolong Diamond (Thermo Fisher) was captured and processed before each imaging session using Zen Black channel alignment tools. The resulting channel alignment was applied to all images from the same imaging session. Images were SIM^2^ processed using 22 (NPM1 and H2AX) or 12 (DAPI) iterations and 0.015 (NPM1 and H2AX) or 0.15 (DAPI) reg weight and were scaled to the raw image. Standard deviation Z projections were made using FIJI. All images are representative of a biological triplicate or greater.

### RNA-Fluorescence in situ hybridization and immunofluorescence

C57BL/6J, testes were collected and fixed in 4% (w/v) PFA for 20 hours at 4°C. Tissues were subsequently rinsed in PBS, dehydrated in ethanol, embedded in paraffin wax, and cut into 4 μm sections. Immunofluorescence was first performed as described above, with the exception of 3% BSA (w/v) in 0.1% tween PBS used as a blocking buffer and quadruple the standard amount of primary antibody. After the washes following secondary antibody incubation, slides were fixed in 4% PFA for 20 minutes at room temperature followed by three 5-minute washes in 0.1% tween PBS all with light protection. HCR^TM^ Gold RNA-FISH (Molecular Instruments) was then performed according to the manufacturer’s instructions using probes that tile the 5’ETS of 47S or that tile the ITS2 sequence. Samples were mounted in DAPI Fluoromount-G (SouthernBiotech) and stored at 4°C with light protection.

### RNA-Fluorescence in situ hybridization and immunocytochemistry

C57BL/6J testes from 60 - 180 dpp animals were collect in PBS and nuclear spreads were prepared as in^68^. Immunocytochemistry was performed first as described in^43^, with the exception of using quadruple the amount of standard primary antibody. After the washes following secondary antibody incubation, slides were fixed in 4% (w/v) PFA for 20 minutes at room temperature followed by three 5-minute washes in 0.1% tween PBS all with light protection. HCR^TM^ Gold RNA-FISH (Molecular Instruments) was then performed according to the manufacturer’s instructions using probes that tile the 5’ETS of 47S or that tile the ITS2 sequence. Samples were mounted in DAPI Fluoromount-G and stored at 4°C with light protection.

### Western blotting

Tissues from C57BL/6J mice aged 5 days post-partum (dpp), 10 dpp, 15 dpp, 18 dpp, 21 dpp, 25 dpp, 28 dpp, and 60 - 180 dpp were collected and flash frozen in liquid nitrogen. Tissue was ground, protein extracted, quantified, and diluted as in^69^. Either 10 µg or 20 µg of total protein was loaded per lane on either a 12% or 15% acrylamide gel. After wet transfer to a PVDF membrane, the membranes were washed in 0.1% PBS tween or 0.1% TBS tween and blocked in 5% nonfat milk. Membranes were incubated overnight at 4°C with the indicated dilution of primary antibody, see Supplementary Table 1. In the case of multiplexing antibodies on the same blot, the concentration of antibody used was increased by a factor of 10. The following day, membranes were washed in 0.1% PBS Tween or 0.1% TBS Tween and incubated with the appropriate Alexa Fluor or horseradish peroxidase (HRP) conjugated secondary antibodies for one hour at room temperature. Membranes were then washed in 0.1% PBS Tween or 0.1% TBS Tween and developed with Pierce ECL Western Blotting Substrate (Thermo Fisher). Membranes were visualized using an Azure Biosystems C600 imager (Azure biosystems). To confirm equal loading and transfer, membranes were stained with Coomassie Brilliant Blue for five minutes at room temperature and subsequently rinsed in ddH_2_O until total protein was visible. Membranes were again visualized using an Azure Biosystems C600 imager. All blots are representative of biological and technical triplicate or greater. For precise experimental details for each primary antibody, see Supplementary Table 1.

### Immunoprecipitation

C57BL/6J testes from 60 - 180 dpp animals were collected and flash frozen with each testes pair split between NPM1 and IgG control IP. Tissue was ground and protein extracted using RIPA (50mM Tris-HCl (pH 8), 150 mM NaCl, 1% (v/v) NP-40 alternative, 0.5% (w/v) sodium deoxycholate, 0.1% (w/v) SDS) supplemented with protease inhibitors (Thermo Scientific, 1 tablet / 10 mL) and RNase inhibitor (NEB, 40 Units / mL)) at a ratio of 1 mL buffer per 100 mg of ground tissue. Protein lysate was then precleared using either Protein A or Protein G beads (Thermo Fisher) that were equilibrated in RIPA (10 μL beads per 100mg tissue) for one hour at 4°C with rotation. For every 1 mL of lysate, either 4 μg of NPM1-L (Proteintech) or mouse IgG antibody (Thermo Fisher) or 2 μg of pan-NPM1 (Proteintech) or rabbit IgG antibody (Thermo Fisher) was added and incubated overnight at 4°C with rotation. The next day beads were added for 2 hours at 4°C with rotation followed by washing with 0.25X TBS (5 mM Tris Base, 37.5 mM NaCl) supplemented with RNase inhibitor (40 Units / mL). Protein was then extracted from the beads by boiling at 95°C for 5 minutes in SB loading dye ((100mM Tris-HCl (pH 6.8), 4% (w/v) SDS, 0.05% (w/v) bromophenol blue, 20% (w/v) glycerol), 50 μL / 1 mL lysate). Eluent along with controls (input, flow-through, and first wash) were prepared for western blotting by dilution in either RIPA buffer (eluent) or SB loading dye (controls) and 1M dithiothreitol (DTT) in a 2:2:1 lysate: RIPA / SB: DTT ratio. Samples were boiled at 95°C for 5 minutes just prior to loading. Western blotting was then performed as described above. Each IP was performed in triplicate.

### RNA extraction and RT-PCR

C57BL/6J testes from 60 - 180 dpp animals were collected and flash frozen prior to RNA extraction. Tissue was ground and RNA was extracted using the miRNeasy Mini Kit (Qiagen) per manufacturer’s instructions. Extracted RNA was DNAse I (Qiagen) treated on-column as per manufacturer’s recommended methods. Final elution was in 30 μL of RNase-free water and RNA was quantified by Nanodrop prior to cDNA synthesis by SuperScript III First-Strand Synthesis System (Thermo Fisher) with random hexamer priming as per manufacturer’s directions. PCR was then performed using PCR Master Mix (2X) (Thermo Fisher) and the primers against specific *Npm1* isoforms described in the Key Resources Table. PCR products were visualized on an agarose gel using an Azure Biosystems C600 imager.

### *NPM1* isoform tissue expression

Adult mouse 100 bp paired end RNA-seq red data were obtained from ENCODE^32^ (www.encodeproject.org) for samples^31^ and accessions as in^70^. Individual samples were aligned to the expanded transcriptome and expression estimated via RSEM. Transcripts per million (TPM) for each of the known *Npm1* isoforms was calculated as the average of available technical replicates.

## Quantification and statistical analysis

RStudio (Posit.co) was used create all graphs and Prism 10 (Graphpad) was used to conduct all statistical analyses. The statistical details of each experiment can be found in the figure legends. Significance was determined via a one-way ANOVA and defined as p 0.05.

### Peason correlation coefficient analysis

SIM images were captured and processed as described above. Biological triplicate was used, with 14-23 cells per stage per animal analyzed (40-72 total cells / stage). Using FIJI, the panNPM1 or the H2AX channel were processed into standard deviation Z projections. Next, either the panNPM1 signal corresponding to the entire XY-body nucleolar structure was outlined by hand and used as ROIs (Fig. 1E) or the H2AX XY-body signal was outlined by hand and used as ROIs (Supp Fig. 2A). Pearson’s R value of each ROI between the panNPM1 and H2AX Z stacks was then determined using the FIJI plugin BIOP JACoP (PTBIOP) using standard inputs.

### Autosomal nucleoli size, number and average pixel intensity quantification

The same SIM Z stacks as described above were additionally used for this analysis. Using FIJI, the panNPM1 and the DAPI channel were processed into standard deviation Z projections. Next, the entire DAPI signal corresponding to each spermatocyte nucleus was outlined by hand and made into a mask. The resulting mask was transformed into a 0–1 pixel intensity scale and then multiplied by the panNPM1 Z stack to isolate spermatocyte panNPM1 signal only. Subsequently, the panNPM1 ROIs corresponding to the entire XY-body nucleolar structure described above were made into a mask and transformed into a 0 - max pixel intensity scale and then subtracted from the spermatocyte panNPM1 signal to isolate autosomal signal only. The resulting image was then thresholded to 32896 and the FIJI plugin MorphLibJ (IJPB) was used to isolate signal greater than 5000 voxels. Nucleoli were counted and volume assessed using the Connect Components Labeling feature with connectivity set to 26. Average pixel intensity of each nucleolus was measured using the built in FIJI plugin 3D suite.

### Fluorescence plot profile analysis

Standard microscopy images were captured and processed as described above. A biological triplicate was used, with 8 – 20 cells per stage per animal analyzed (30 - 39 total cells / stage). Using FIJI, a 25-pixel length line was drawn from the top of each XY-body associated nucleolar foci through the center. Individual plot profiles were generated for each cell and for all channels using the built-in FIJI tool. Underlying plot data was then exported into Excel and the signal intensity for each pixel distance was averaged across all cells within a developmental stage.

## Additional resources

None

---

**Supp Fig 1.**
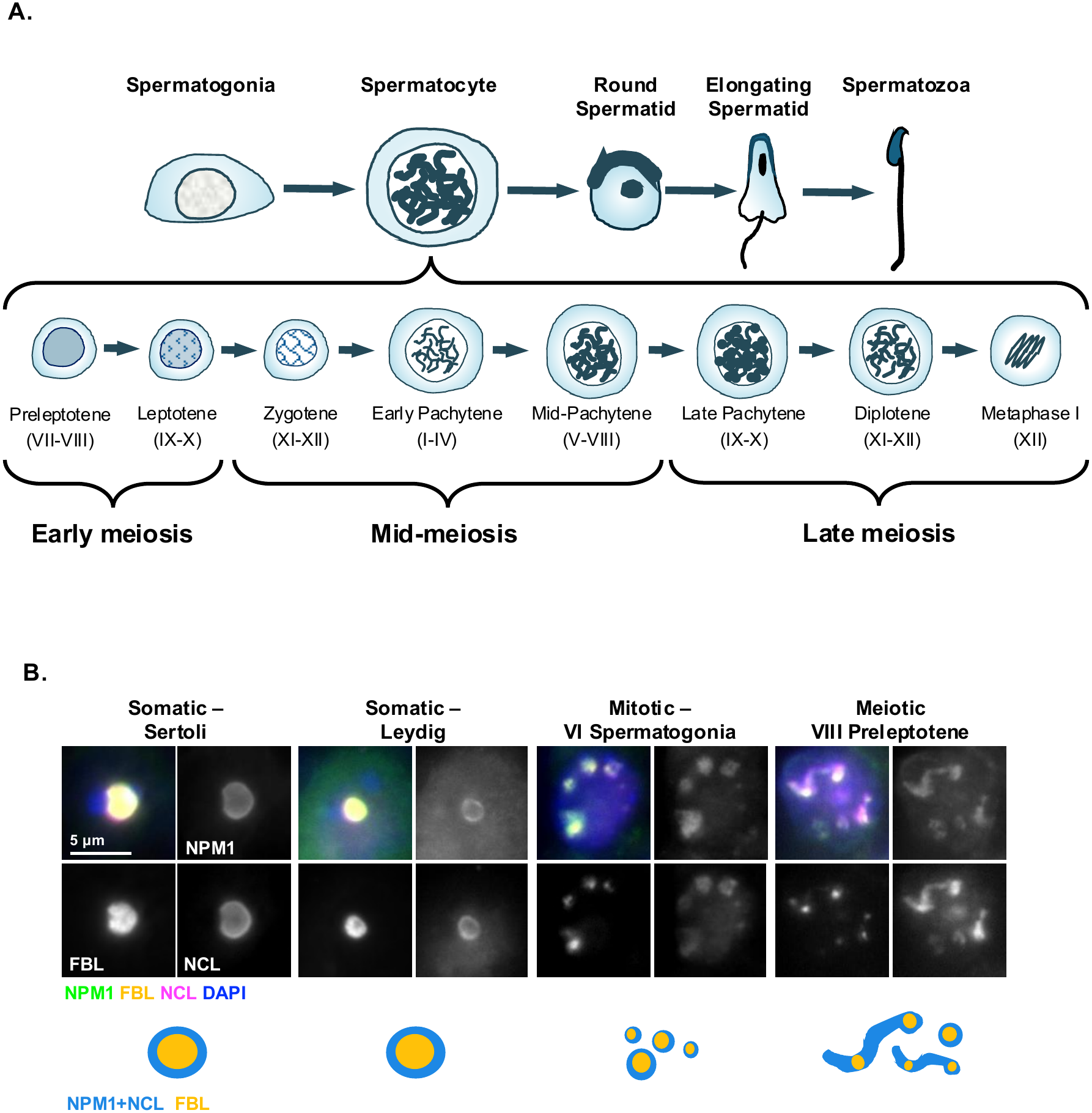
Somatic and mitotic cells of the testis contain morphologically normal nucleoli. **(A)** Schematic of mammalian male germ cells by developmental stage with a focus on meiotic spermatocytes, of which morphological and molecular markers can discern up to seventeen distinct development stages. Derived from figure found in^71^. **(B)** Immunofluorescence and schematic of nucleolar markers NPM1, FBL, and NCL in adult testis cell types with a focus on somatic, mitotic, and early meiotic cells. Meiotic substage indicated with stage of the cycle of the seminiferous epithelium designated by Roman numeral. Images are representative of biological triplicate.

**Supp Fig 2.**
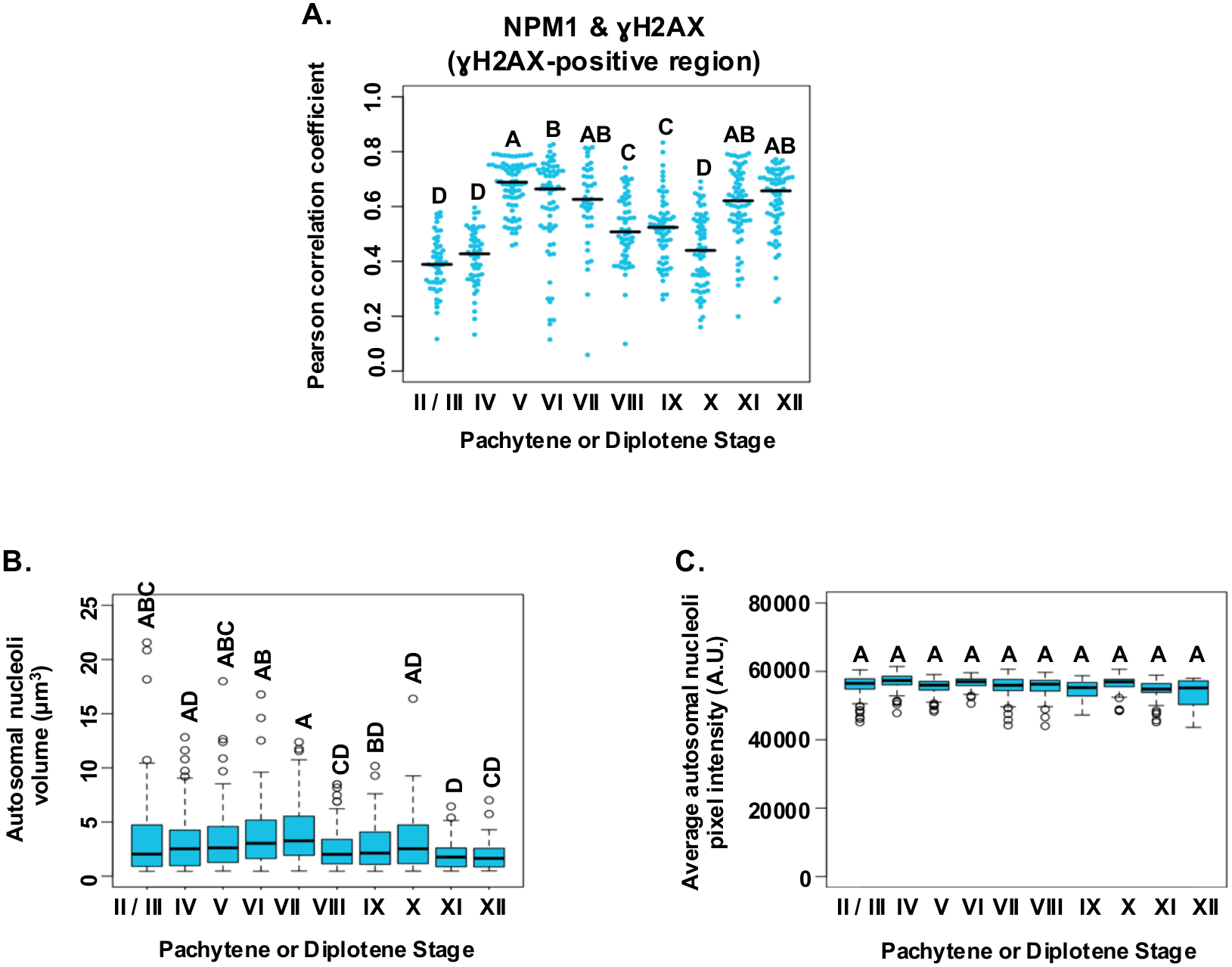
NPM1 and H2AX colocalization is linked to meiotic subphase. **(A)** Colocalization coefficients of NPM1 and□H2AX in the XY-body by developmental phase of meiotic spermatocytes. Dots are individual cells. Quantification of **(B)** non XY-body associated NPM1 foci volume and **(C)** the average pixel intensity of non XY-body associated NPM1 foci by pachytene or diplotene stage. Letters indicate significance levels as determined by one-way ANOVA (p < 0.05). Bar – sample median. Values reported for individual cells from biological triplicate (14 – 23 cells / stage / animal). Roman numerals indicate spermatogenic stage. See Star Methods for all image processing and quantification details.

**Supp Fig. 3.**
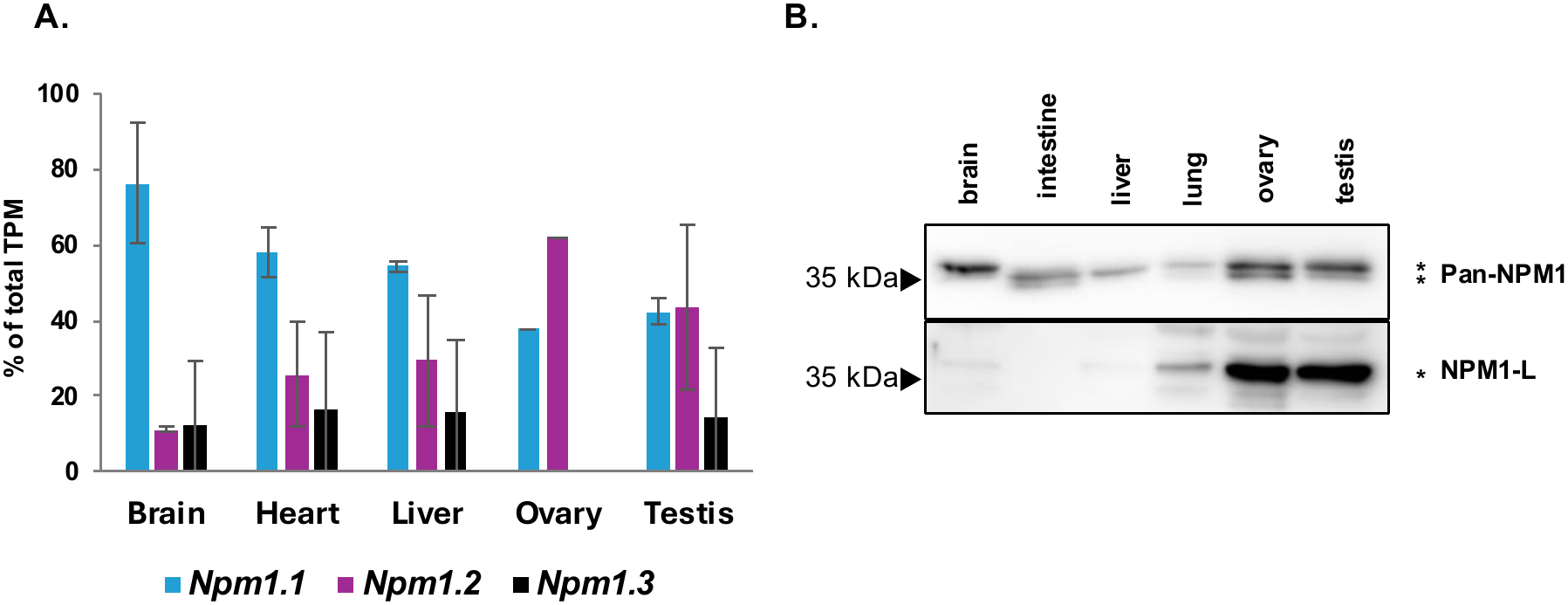
A short isoform of NPM1 is enriched in reproductive tissues. **(A)** Tissue expression levels of *Npm1* isoforms as a percentage of total *Npm1* expression by RNA-sequencing. Error bars - standard deviation. **(B)** Western blot of panNPM1 and NPM1-L antibodies on adult tissues. See Supp Fig. 7 for loading control. Blots are representative of biological triplicate.

**Supp Fig. 4.**
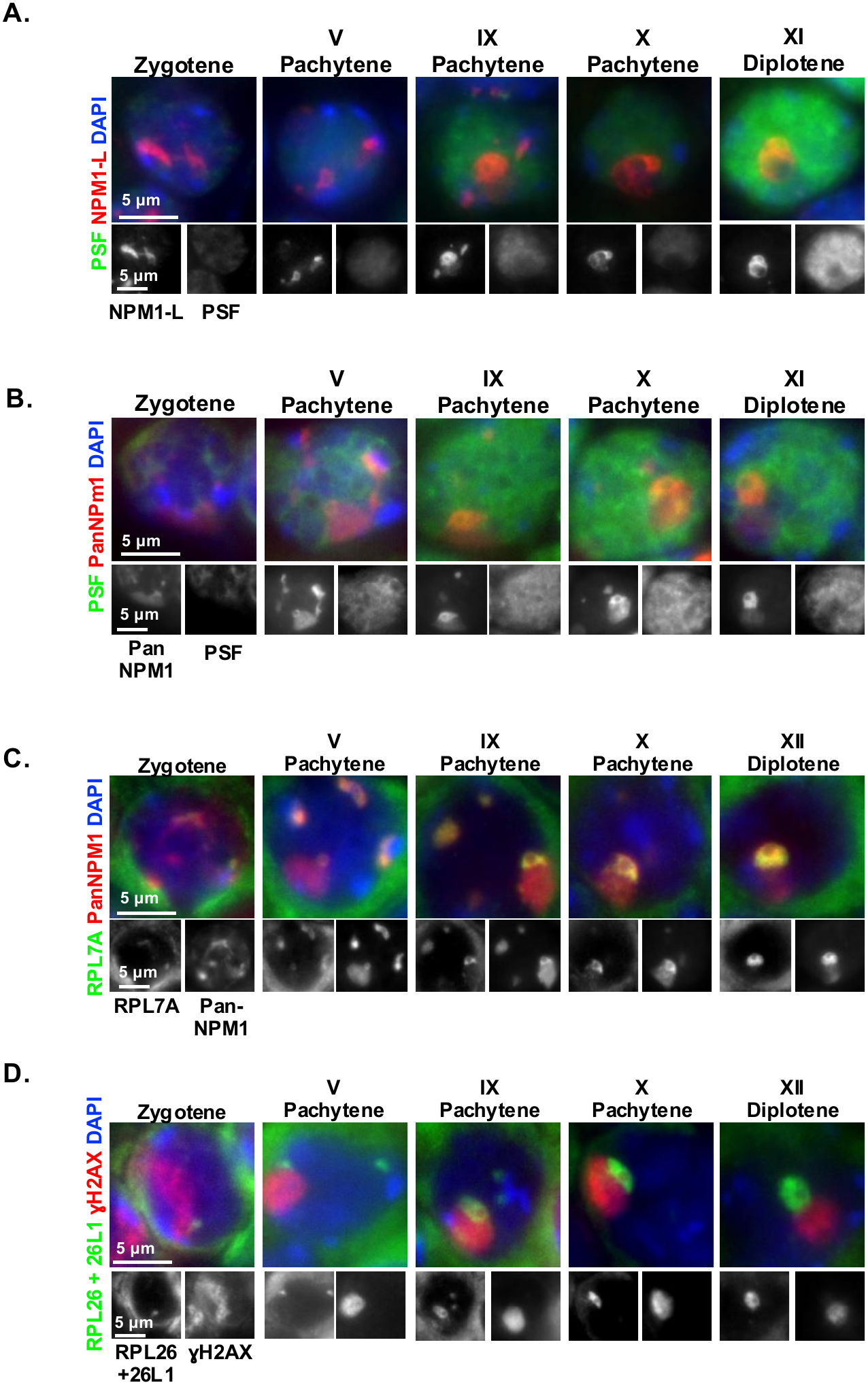
All NPM1-positive bodies of meiotic male germ cells contain markers indicative of active nucleoli. Immunofluorescence of PSF and **(A)** NPM1-L or **(B)** panNPM1 in meiotic spermatocytes by developmental phase. Immunofluorescence of panNPM1 and **(C)** RPL7A or **(D)** RPL26 / 26L1 in meiotic spermatocytes by developmental phase. Meiotic substage indicated with stage of the cycle of the seminiferous epithelium designated by Roman numeral. Images are representative of biological triplicate.

**Supp Fig. 5.**
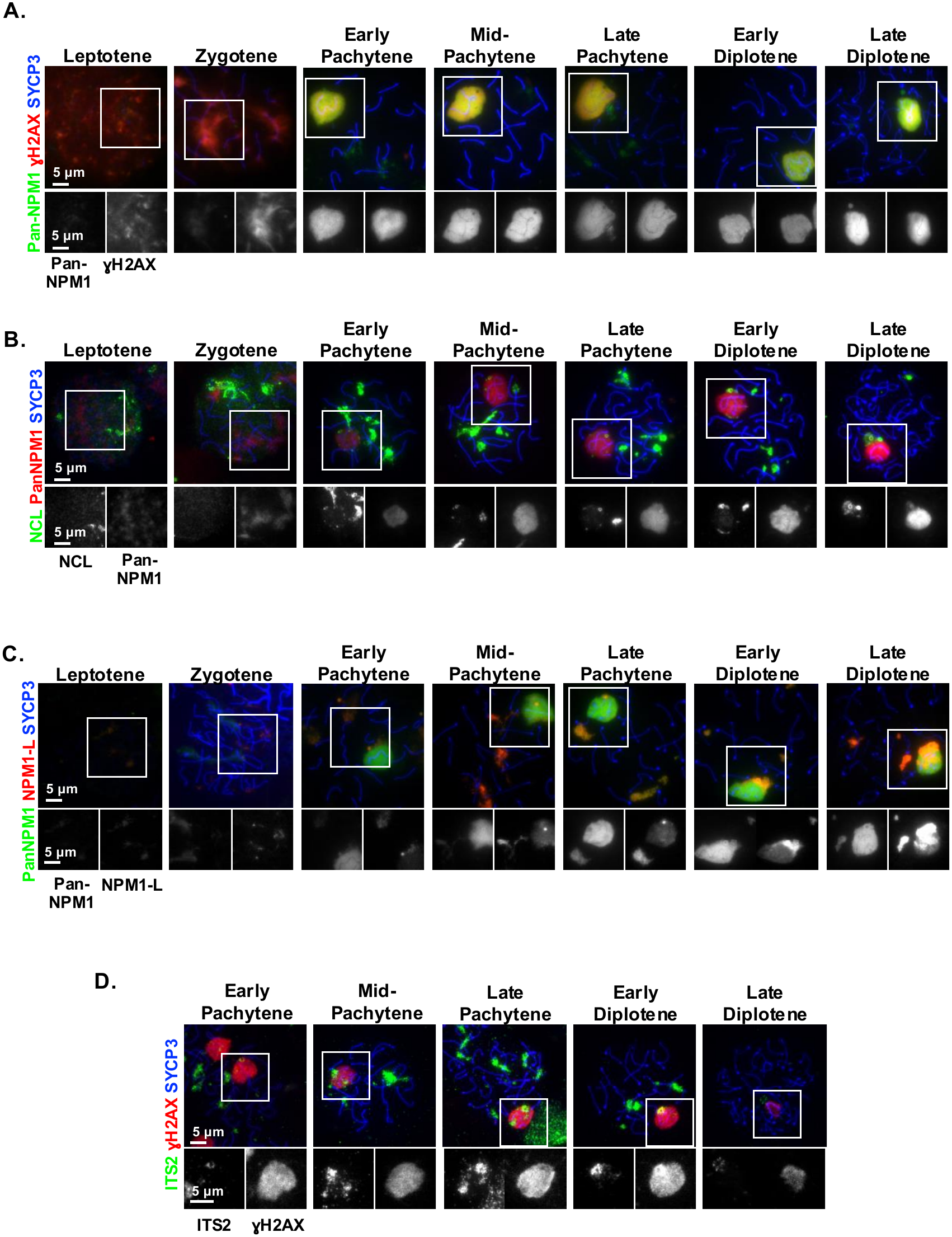
The separation of NPM1-only and XY-body adjacent NFN bodies is altered in nuclear spreads. Immunocytochemistry on spermatocyte nuclear spreads of **(A)** panNPM1,□H2AX, and the meiotic spermatocyte stage marker SYCP3; **(B)** NCL, panNPM1, and SYCP3; and **(C)** panNPM1, NPM1-L, and SYCP3 by meiotic substage. **(D)** RNA-FISH of ITS2 pre-rRNA with immunocytochemistry of SYCP3 and□H2AX. Pixel intensity scale matched across images except for increased panNPM1 channel in leptotene and zygotene stages for visibility. Images are representative of biological triplicate.

**Supp Fig. 6.**
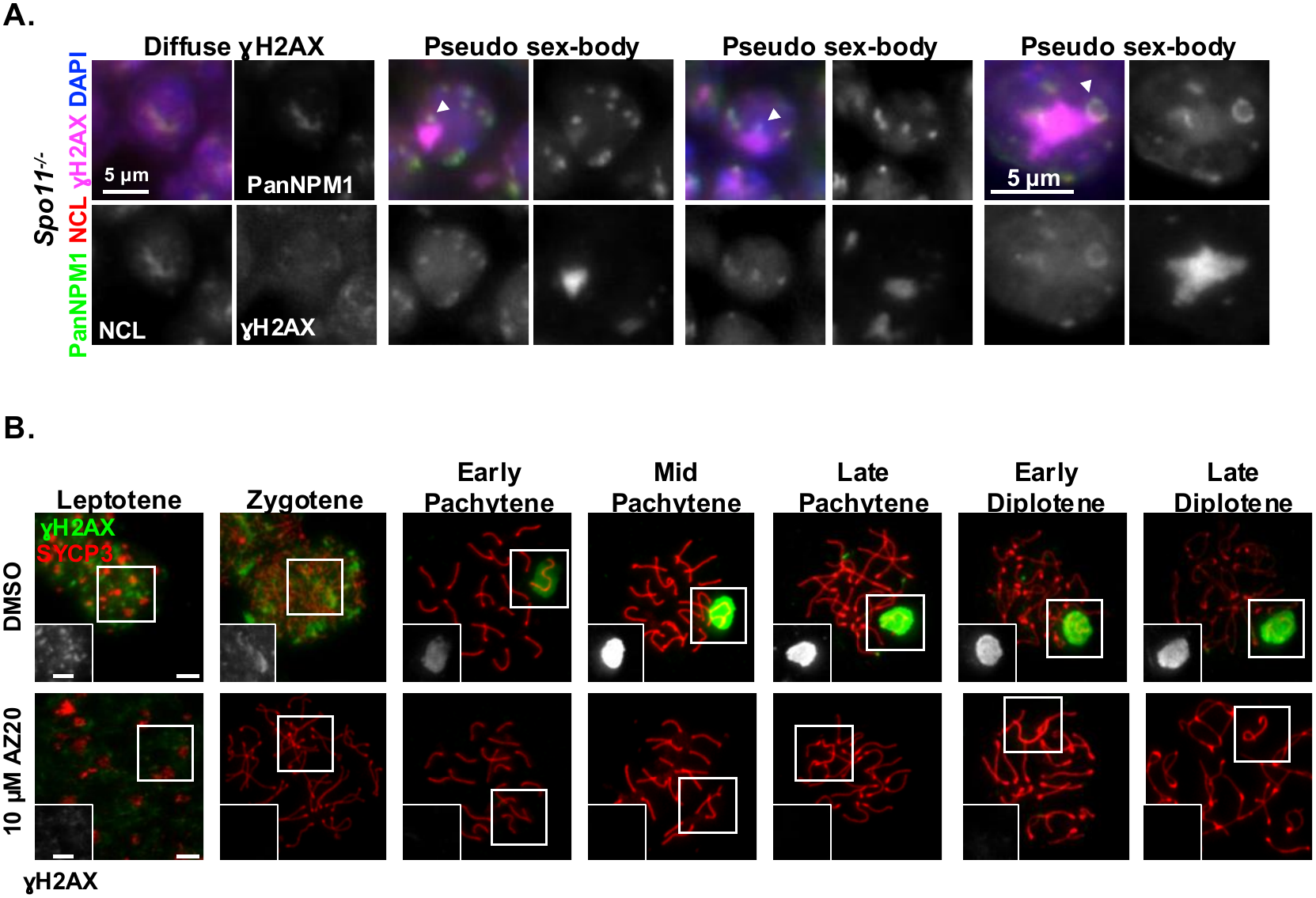
Meiotic nucleoli are reliant on formation of a H2AX domain. **(A)** Immunofluorescence of panNPM1, NCL, and□H2AX in adult *Spo11* mutant spermatocytes with or without a pseudo sex-bodies. Arrow – pseudo sex-body adjacent NFN body. **(B)** Immunocytochemistry of H2AX (inset) and SYCP3 on cultured spermatocytes. Scale - 5 μm. Images are representative of biological triplicate.

**Supp. Fig. 7.**
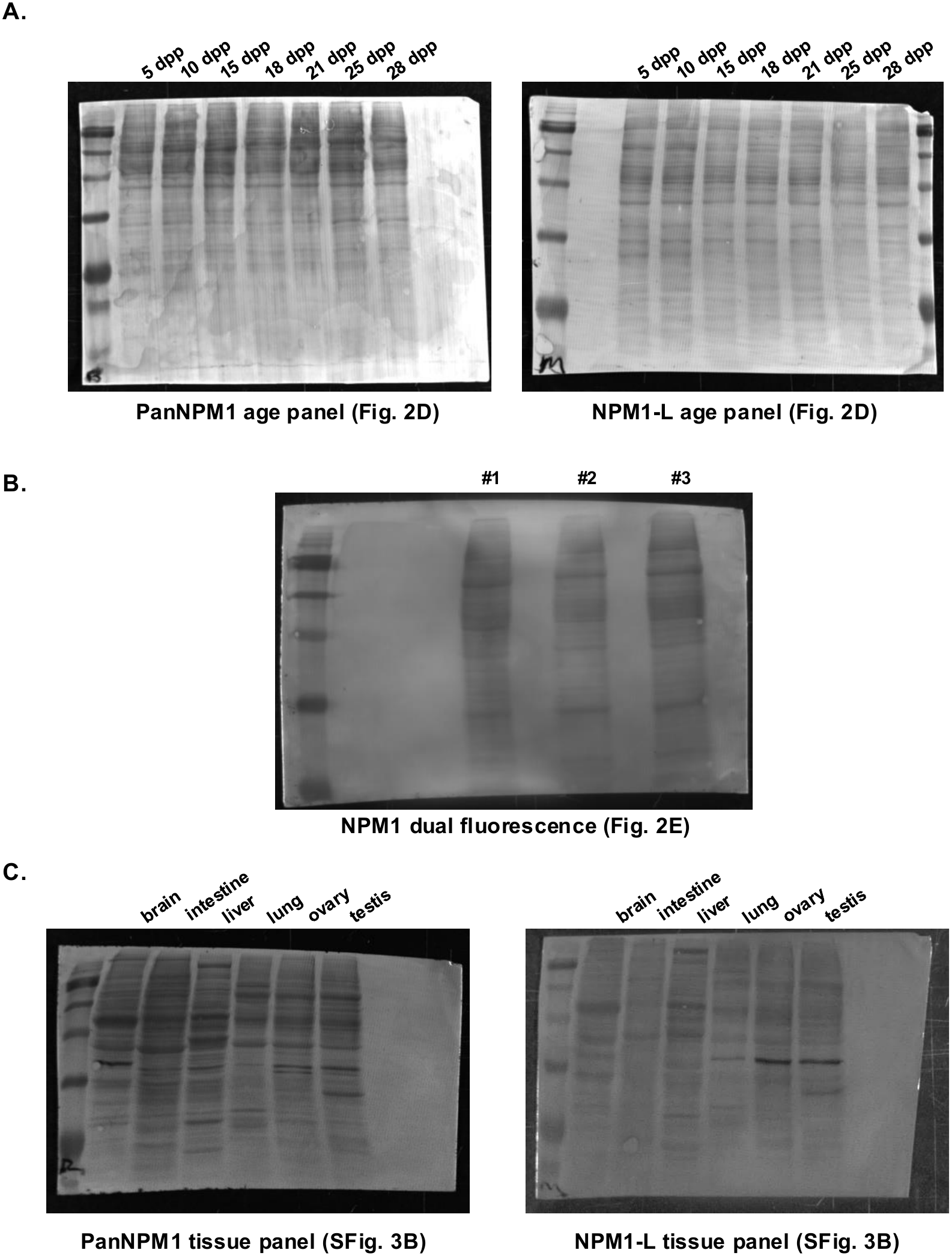
Western blot loading controls. Coomassie blue loading and transfer control for western blots in **(A)** Fig. 2D, **(B)** Fig. 2E, and **(C)** Supp Fig. 3B.

**Supp. Fig 8.**
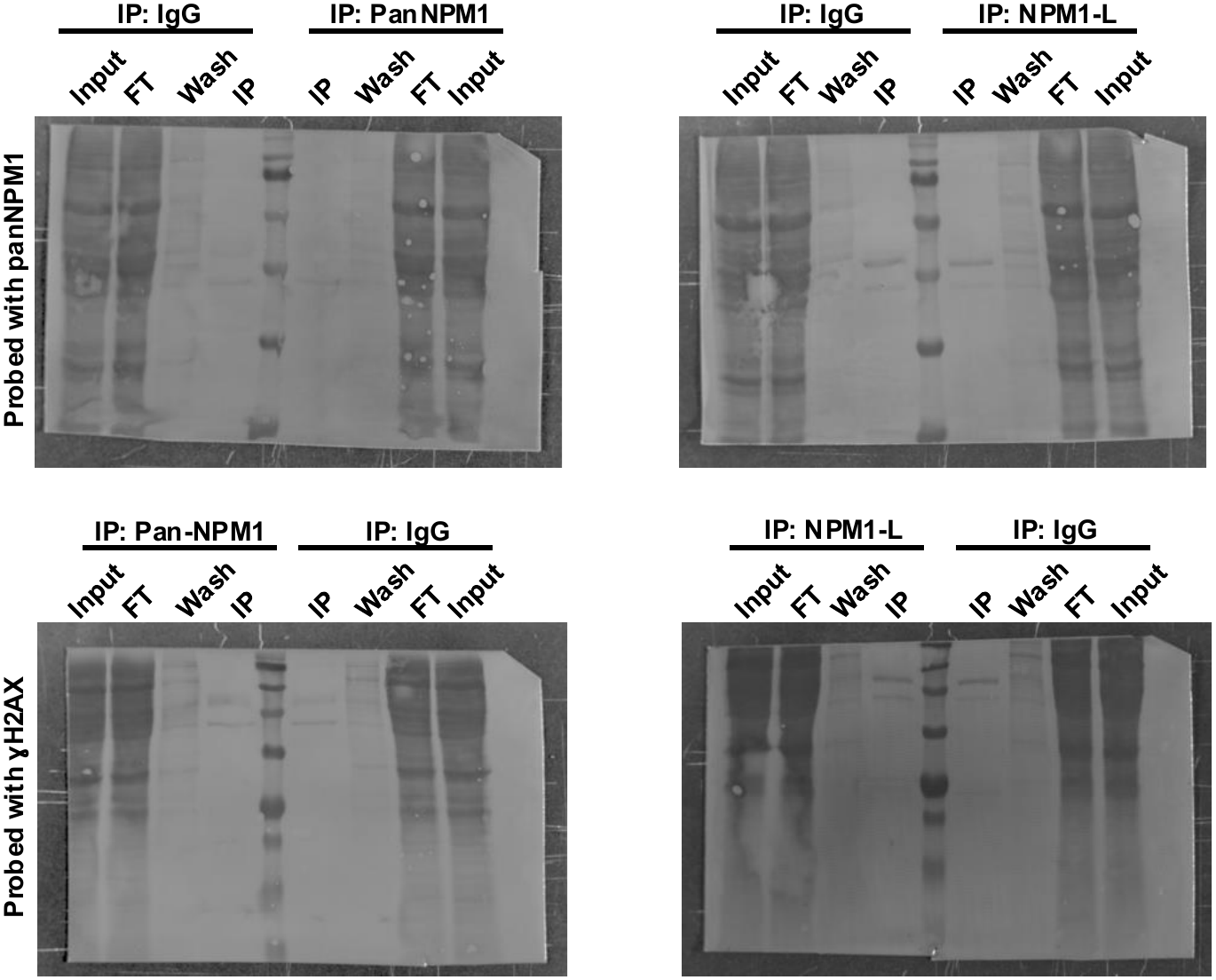
Western blot loading controls for immunoprecipitation. Coomassie blue loading and transfer control for immunoprecipitation blots in Fig. 4.

