## Supplementary material for "A short nucleophosmin isoform couples nucleolar remodeling to DNA damage repair during meiosis": RNA-FISH Probe Sequences

**47S: 20 probes tiling the following 5’ ETS sequence** Tactgacacgctgtcctttccctattaacactaaaggacactataaagagaccctttcgatttaaggctgttttgcttgtccagcctattctttttactggcttgggtctgtcgcggtgcctgaagctgtccccgagccacgcttcctgctttcccgggcttgctgcttgcgtgtgcttgctgtgggcagcttgtgacaactgggcgctgtgactttgctgcgtgtcagacgtttttcccgatttccccgaggtgtcgttgtcacacctgtcccggttggaatggtggagccagctgtggttgagggccaccttatttcggctcactttttttttttttttttctcttggagtcccgaacctccgctcttttctcttcccggtctttcttccacatgcctcccgagtgcatttctttttgttttttttctttttttttttttttttttggggaggtggagagtcccgagtacttcactcctgtctgtggtgtccaagtgttcatgccacgtgcctcccgagtgcacttttttttgtggcagtcgctcgttgtgttctcttgttctgtgtctgcccgtatcagtaactgtcttgccccgcgtgtaagacattcctatctcgcttgtttctcccgattgcgcgtcgttgct

**ITS2: 20 probes tiling the following sequence**

cagacgtgtgggcggttgtcggtgtggcgcgcgcgcccgcgtcgcggagcctggtctcccccgcgcatccgcgctcgcggcttcttcccgctccgccgttcccgccctcgcccgtgcaccccggtcctggcctcgcgtcggcgcctcccggaccgctgcctcaccagtctttctcggtcccgtgccccgtgggaacccaccgcgcccccgtggcgcccgggggtgggcgcgtccgcatctgctctggtcgaggttggcggttgagggtgtgcgtgcgccgaggtggtggtcggtcccctgcggccgcggggttgtcggggtggcggtcgacgagggccggtcggtcgcctgcggtggttgtctgtgtgtgtttgggtcttgcgctgggggaggcggggtcgaccgctcgcggggttggcgcggtcgcccggcgccgcgcaccctccggcttgtgtggagggagagcgagggcgagaacggagagaggtggtatccccggtggcgttgcgagggagggtttggcgtcccgcgtccgtccgtccctccctccctcggtgggcgccttcgcgccgcacgcggccgctaggggcggtcggggcccgtggcccccgtggctcttcttcgtctccgcttctccttcacccgggcggtacccgctccggcgccggcccgcgggacgccgcggcgtccgtgcgccgatgcgagtcacccccgggtgttgcgagttcggggagggagagggcctcgctgacccgttgcgtcccggcttccctgggggggacccggcgtctgtgggctgtgcgtcccgggggttgcgtgtgagtaagatcctccacccccgccgccctcccctcccgccggcctctcggggaccccctgagacggttcgccggctcgtcctcccgtgccgccgggtgccgtctctttcccgcccgcctcctcgctctcttcttcccgcggctgggcgcgtgtccccccttt
