## Supplementary Table 1 for "A short nucleophosmin isoform couples nucleolar remodeling to DNA damage repair during meiosis"

| Antibody | IF conditions | ICC conditions | FISH / IF conditions |
|---|---|---|---|
| Fibrillarin | 1:100 | N.A. | N.A. |
| Rabbit NPM1 Proteintech | 1:1000 | 1:1000 | 1:250 |
| Mouse NPM1 Proteintech | 1:400 | N.A. | N.A. |
| Mouse NPM1 Abcam | 1:1000 | 1:1000 | N.A. |
| Nucleolin | 1:500 | 1:500 | N.A. |
| Mouse $\gamma$H2AX | 1:1000 | N.A. | 1:250 |
| PSF | 1:500 | N.A. | N.A. |
| RPL7A | 1:1000 | N.A. | N.A. |
| RPL26/26L1 | 1:500 | N.A. | N.A. |
| Pseuodouridine | 1:200 | N.A. | N.A. |
| Rabbit $\gamma$H2AX | N.A. | 1:3000 | N.A. |
| SYCP3 | N.A. | 1:200 | 1:100 |
| BRCA1 | N.A. | 1:500 | N.A. |
| Goat anti-Mouse Alexa Fluor™ 488 | 1:1000 | 1:1000 | N.A. |
| Goat anti-Mouse Alexa Fluor™ 594 | 1:1000 | 1:1000 | N.A. |
| Goat anti-Mouse Alexa Fluor™ 680 | 1:1000 | 1:1000 | 1:1000 |
| Goat anti-Rabbit Alexa Fluor™ 488 | 1:1000 | 1:1000 | N.A. |
| Goat anti-Rabbit Alexa Fluor™ 594 | 1:1000 | 1:1000 | 1:1000 |
| Goat anti-Rabbit Alexa Fluor™ 680 | 1:1000 | 1:1000 | N.A. |
| Goat Anti-Rabbit IgG (H+L) -HRP Conjugate | N.A. | N.A. | N.A. |
| Goat Anti-Mouse IgG (H+L) -HRP Conjugate | N.A. | N.A. | N.A. |

| Western blot conditions |
| --- |
| N.A. |
| 1:4000, 15% gel |
| 1:1000, 15% gel |
| N.A. |
| N.A. |
| N.A. |
| N.A. |
| N.A. |
| N.A. |
| N.A. |
| 1:1000 |
| N.A. |
| N.A. |
| 1:4000 |
| N.A. |
| N.A. |
| 1:1000 |
| 1:4000 |
| N.A. |
| 1:2000 |
| 1:2000 |
